# Comparing multifunctional viral and eukaryotic proteins for generating scission necks in membranes

**DOI:** 10.1101/2024.01.05.574447

**Authors:** Haleh Alimohamadi, Elizabeth Wei-Chia Luo, Shivam Gupta, Jaime de Anda, Rena Yang, Taraknath Mandal, Gerard C. L. Wong

**Affiliations:** Department of Bioengineering, University of California, Los Angeles, Los Angeles, CA 90025, USA; Department of Chemistry and Biochemistry, University of California, Los Angeles, CA, 90095, USA; Department of Microbiology, Immunology, and Molecular Genetics, University of California, Los Angeles, CA, 90095, USA; California NanoSystems Institute, University of California, Los Angeles, CA, 90095, USA; Department of Physics, Indian Institute of Technology Kanpur, Kanpur, 208016, India

## Abstract

Deterministic formation of membrane scission necks by protein machinery with multiplexed functions is critical in biology. A microbial example is the M2 viroporin, a proton pump from the influenza A virus which is multiplexed with membrane remodeling activity to induce budding and scission in the host membrane during viral maturation. In comparison, the dynamin family constitutes a class of eukaryotic proteins implicated in mitochondrial fission, as well as various budding and endocytosis pathways. In the case of Dnm1, the mitochondrial fission protein in yeast, the membrane remodeling activity is multiplexed with mechanoenzyme activity to create fission necks. It is not clear why these functions are combined in these scission processes, which occur in drastically different compositions and solution conditions. In general, direct experimental access to changing neck sizes induced by individual proteins or peptide fragments is challenging due to the nanoscale dimensions and influence of thermal fluctuations. Here, we use a mechanical model to estimate the size of scission necks by leveraging Small-Angle X-ray Scattering (SAXS) structural data of protein-lipid systems under different conditions. The influence of interfacial tension, lipid composition, and membrane budding morphology on the size of the induced scission necks is systematically investigated using our data and molecular dynamic simulations. We find that the M2 budding protein from the influenza A virus has robust pH-dependent membrane activity that induces nanoscopic necks within the range of spontaneous hemi-fission for a broad range of lipid compositions. In contrast, the sizes of scission necks generated by mitochondrial fission proteins strongly depend on lipid composition, which suggests a role for mechanical constriction.

## Introduction

Membrane neck formation is a critical step in many eukaryotic transport processes such as endocytosis, exocytosis, and intracellular trafficking ^1–3^. For microbial processes, vesicle budding and membrane neck formation also play crucial roles in the replication cycle of enveloped viruses and parasites ^4–6^. In both classes of examples, the protein machinery often multiplexes membrane remodeling activity with drastically different functions. The formation of scission necks in eukaryotes is controlled by proteins that adsorb or insert into the cell membrane and reshape the membrane into distinct morphologies via different functions ^7–9^. For example, dynamin-related GTPase proteins (Dnm1 in yeast, Drp1 in mammals) exhibit mechanoenzyme activity to ‘squeeze’ fission necks, but also the ability to remodel membrane curvature, thereby facilitating the formation of membrane necks in mitochondrial fission in a process that is not completely understood ^10–14^. Similarly, bacterial pathogens and viruses including Rabies virus, Parainfluenza virus-5, and influenza A possess protein machinery capable of membrane remodeling to enforce budding and scission in host cell membranes ^4–6^. However, this activity is also often combined with other functions. In the case of Influenza, A, the M2 viroporin that mediates budding is also a proton pump that can acidify local environments. Presently, these multiplexed functions are considered serendipitous accidents in evolution. There is no conceptual framework to see why these functions in particular are combined or whether the additional functions are somehow synergistic with membrane remodeling.

To understand the physicochemical principles underlying the membrane remodeling required for scission pores, it is important to experimentally determine the size and geometry of scission necks across a diverse range of experimental conditions, given that neck formation is distinct for different biological contexts (local lipid compositions) and local environmental conditions (pH, temperature). Various techniques including X-ray crystallography ^15^, nuclear magnetic resonance spectroscopy (NMR)^16^, and electron microscopy (EM) ^17^ can in principle be used for estimating membrane curvatures in scission necks of reconstituted systems. However, these techniques are usually applicable only to specific experimental conditions that are amenable to the measurement itself (such as freezing or vacuum conditions). Moreover, measuring membrane neck sizes is further complicated by the dynamics of membrane deformations, the nanoscopic scale of neck formation, and the heterogeneous, transient, fluctuating nature of such structures^18^.

Small-Angle X-ray Scattering (SAXS) ^19,20^ is a non-destructive technique that has been used to study the structure and phase behavior of biomolecular systems ^21^. Specifically, using SAXS, previous studies have shown that pore-forming proteins can reorganize small unilamellar vesicles (SUVs) into bicontinuous cubic phases ^22–24^. Cubic phases are periodic minimal surfaces that have been found in a wide range of membrane-bound organelles such as endoplasmic reticulum, Golgi apparatus, and mitochondrial membranes ^25–28^. Cubic phases are idealized surfaces with zero mean curvature but rich in bilayer negative Gaussian curvature (NGC) (Fig. 1). NGC is a geometric requirement of scission neck formation, so the observation of NGC-rich phases is consistent with expectation (Fig. 1). However, the precise quantitative relationship between the extent of induced curvatures observed in cubic phases (for instance, the NGC density per volume) and the actual dimensions of scission necks is not clear (Fig. 1). The ability to estimate scission neck dimensions from the structure of cubic phases can have great utility since SAXS can be used for different lipid compositions and for different intensive thermodynamic variables relevant to different biological environments.

**Figure 1.**
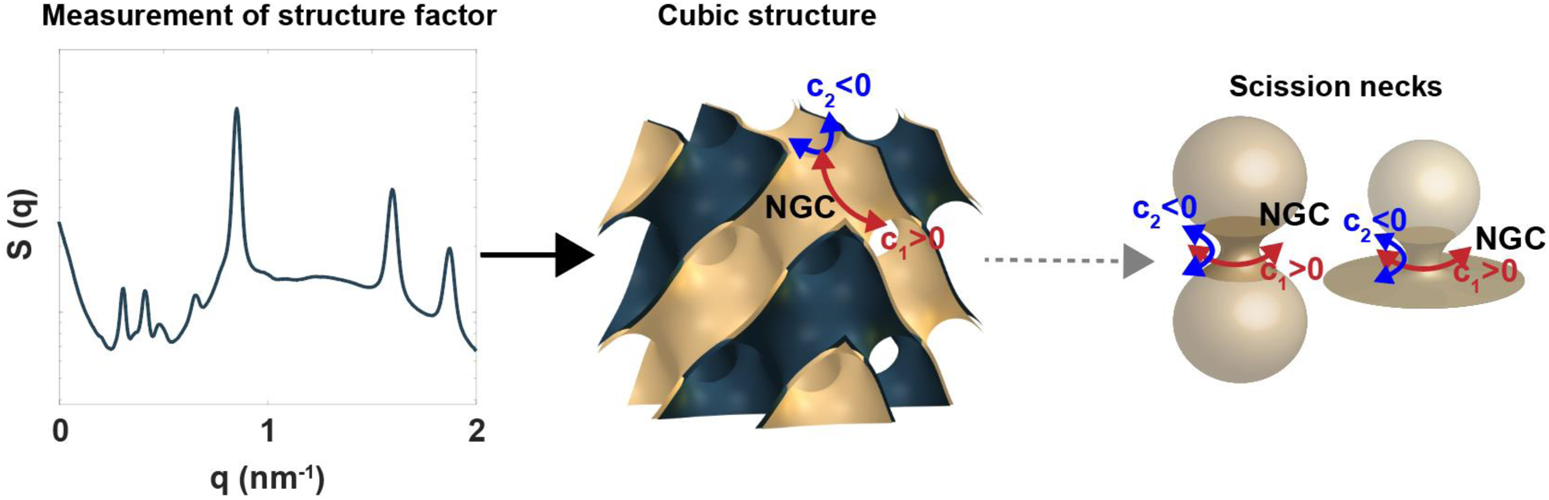
Estimating the induced radius of scission necks by membrane-remodeling proteins based on the membrane deformations observed in SAXS spectra. Cubic structures and catenoid-shaped scission necks are idealized minimal surfaces with zero mean curvature and negative Gaussian curvature (c_1_ > 0 and c_2_ < 0).

The use of theoretical and computational methods integrated with experimental information has emerged as a potent approach for investigating induced membrane curvatures by proteins ^9,29–33^, and for distilling insights that are otherwise challenging to obtain. For example, Kozlovsky et al. have studied the role of proteins in membrane neck formation during fission and fusion processes ^30,34^. They showed that for wide necks (neck radius > monolayer thickness), the necks adopt the shape of a catenoid with zero mean curvature ^30^. Chabanon et al. proposed that the Gaussian curvature of neck-like structures governs the distribution of the proteins in the neck region ^31^. Dharmavaram et al. suggested that the mismatch between the Gaussian curvature of the partially assembled capsid as a spherical cap and the catenoid-shaped neck region creates an energy barrier during viral budding ^29^.

In this study, we construct a minimal continuum elastic membrane model, and use it to quantitatively extract protein-induced NGC from structures of cubic phases measured in SAXS. We apply the derived induced curvature as an input to the free energy of the membrane neck formation and estimate the radius of a stable scission neck via energy minimization. We introduce a dimensionless quantity, the “spontaneous scission number (SSN)” as a function of the cubic lattice constant, protein size, and size of the neck. The SSN controls the degree of the scission neck constriction, and we found that SSN > 1 is a threshold for the formation of necks sufficiently narrow to be within the range of possible spontaneous hemi-fission.

We apply our mechanical framework to estimate the size of scission necks induced by mitochondrial fission proteins in yeast (Dnm1) and the M2 budding protein from the influenza A virus, using measured cubic lattice constants ^22,23^. We show that the degree of neck constriction by mitochondrial fission protein Dnm1 depends highly on lipid composition. Not only is this consistent with expectations for a highly regulated cellular process, but it also suggests that the mechanoenzyme activity may be necessary to supplement the membrane remodeling activity when lipid compositions are not optimal for membrane remodeling alone. In contrast, the M2 protein from influenza A infection is able to squeeze the budding neck to narrow sizes within the range of spontaneous hemi-fission in a manner that is significantly less sensitive to lipid composition. This is consistent with the need for the virus to successfully mature and bud in a broad range of eukaryotic cells with different lipid compositions. We demonstrate that a synthesized peptide of M2 protein, consisting of the transmembrane helix and the C-terminal cytoplasmic helix (M2TM-cyto), is sufficient to constrict vesicles and form narrow necks. Furthermore, we show that acidification in fact enhances the membrane remodeling activity of the M2 protein, which aligns with the activation of M2 channel proton transport upon acidification ^35^. Using coarse-grained molecular dynamic simulations, we also illustrate that the M2 protein from influenza A virus can induce large anisotropic curvature independent of lipid composition. From these results, we believe our framework can be generalized to other protein-membrane systems.

## Model development

### Membrane mechanics

We model the lipid bilayer as a thin elastic shell with negligible thickness compared to the radii of membrane curvature ^36,37^. We assume the membrane is incompressible since the energetic cost of stretching the membrane is much higher than membrane bending ^35^. We also assume that NGC-generating inclusions are confined in the neck region based on previous studies that have shown that fission and viral proteins such as Dnm1 and M2 can sense the membrane curvature and entropic forces localize them at the scission necks in order to facilitate the constriction process ^6,38,39^. Furthermore, we assume that membrane inclusions are more rigid than the lipid bilayer and use the augmented version of the Helfrich–Canham energy to model the membrane-protein interactions ^40^. Assuming the system is at mechanical equilibrium at all times, the total energy of the system (*E*) is given by ^41–46^

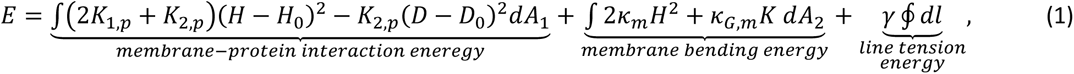

where the first term corresponds to the mismatch elastic energy between the local shape of the lipid membrane and the intrinsic shape of the inclusion in the neck region, the second term indicates the membrane bending energy in the domain with no inclusion, and the third term represents the energy cost due to the phase separation of NGC-generating inclusions at the scission necks. In Eq. 1, *K*_1,*p*_ and *K*_2,*p*_ are constants (*K*_1,*p*_ > 0 and *K*_2,*p*_ < 0) representing the strength of interaction between the inclusions and the surrounding membrane ^41,42^. *K_m_* and *K_G,m_* are the bending and Gaussian moduli of the lipid bilayer, respectively. *H* is the mean curvature, *D* is the curvature deviator, and *K* is the Gaussian curvature of the lipid membrane. *H*_0_ is the spontaneous mean curvature, *D*_0_ is the intrinsic deviatoric curvature, and *γ* is line tension at the neck boundary. In Eq. 1, we assume that the outer surface area of the neck is fully covered by NGC-generating proteins and the first integral is taken over the area of the membrane that is covered by NGC-generating inclusions (*dA*_1_), while the second integral is taken over the bare lipid bilayer with no inclusion (*dA*_2_).

### Free energy of scission neck formation

To establish a relationship between the induced curvatures by membrane-remodeling proteins such as Drp1 observed in SAXS to the actual size of scission necks, we assume that proteins remodel a spherical vesicle with a radius of *R_S_* = 100 nm (approximate size of a SUV in SAXS) to an idealized geometry of mitochondrial fission. The idealized geometry of membrane fission includes a rotationally symmetric catenoid surface with a neck radius of *r_n_* and two spherical caps with a radius of *R* (Fig. 2A). This simplified geometry has been widely used in previous studies to model local neck structures, viral budding, and membrane fission ^29–32^. The excess free energy of the system (including membrane and proteins) with respect to the vesicle can be written as

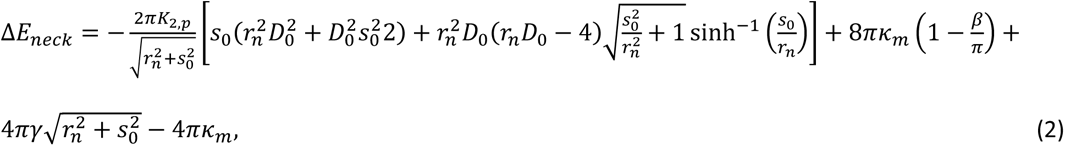

where *κ_m_* is the membrane bending rigidity, *β* is the angle between the vertical axis and edge of the disk forming at the base of the catenoid, and *S*_0_ is the length of the catenoid (Fig. 2A). Here, we assume that the line tension *γ* accounts for the energy cost arising from protein segregation at the scission necks and also the discontinuity in the membrane curvature of the idealized geometry, where the catenoids connect to the spherical cap (analogous to the cup-like model ^44^). It should be mentioned that in Eq. 2, for simplicity, we set *κ_G,m_* (membrane Gaussian modulus) ~ – *κ_m_* and assume that the induced deviatoric curvature by membrane-remodeling proteins is dominant (*H*_0_ ≪ *D*_0_).

**Figure 2.**
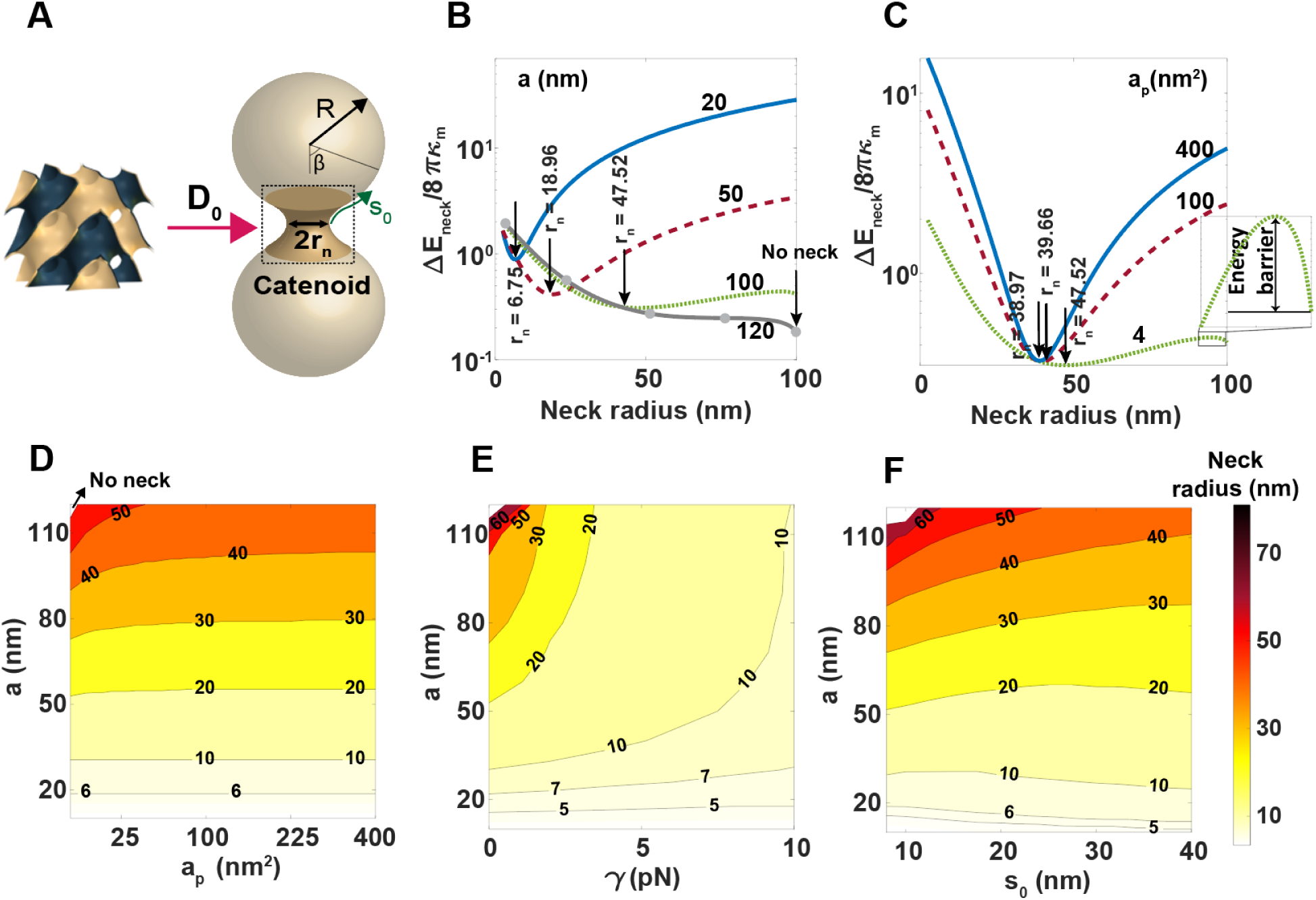
Estimating the radius of induced scission necks by membrane-remodeling proteins using the cubic structures in SAXS. **(A, left)** The induced deviatoric curvature by scission proteins (*D*_0_) is extracted from the structures of cubic phases in SAXS measurements (Eq. 6). **(A, right)** An idealized geometry of a membrane scission neck was modeled as a rotationally symmetric catenoid surface with a neck radius of *r_n_* and two identical spherical caps with a radius of R. **(B)** The change in the energy of the system from a spherical vesicle to the idealized membrane scission neck geometry (Eq. 4) as the function of the neck radius for four different lattice constants (*a_p_* = 4 nm^2^, *S*_0_ = 10 nm and *γ* = 0). Arrows show the location of minimum energy corresponding to the stable radius of the constricted neck. The neck radius increases with the increasing lattice constant of cubic structures. (**C**) The change in the energy of the system as the function of the neck radius for three different protein surface areas in contact with the membrane at fixed *a* = 100 nm (*S*_0_ = 10 nm and *γ* = 0). Larger proteins lead to a smaller scission neck, while at a small protein size, an energy barrier is associated with membrane neck constriction. Contour plot of the membrane scission neck radius for a range of (**D**) cubic lattice constants and protein surface area (*S*_0_ = 10 nm and *γ* = 0), (**E**) cubic lattice constants and line tension (*a_p_* = 4 nm^2^ and *S*_0_ = 10), and (**F**) cubic lattice constants and catenoid length (*a_p_* = 4 nm^2^ and *γ* = 0). The white domains in panels (**D**), (**E**), and (**F**) mark the regions with no energy minima and no neck formation.

The magnitude of induced deviatoric curvature can be estimated from the cubic structure formed by membrane-remodeling proteins in SAXS, given as ^47^

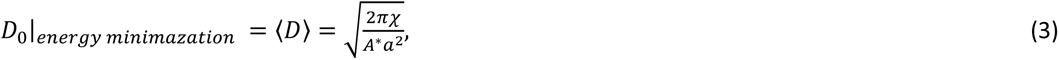

where *a* is the lattice parameter, *χ* is the Euler characteristic, and *A*\* is the surface area per unit cell specific to each cubic phase ^48,49^. Having the amount of induced deviatoric curvature from SAXS data (Eq. 3), we numerically calculate the excess energy of neck formation (Eq. 2) and find the neck radius (*r_n_*) that minimizes *ΔE_neck_*. In our calculations, we estimated the strength of interaction between the membrane and protein (*K*_2,*p*_) based on the elastic properties of the membrane and the size of the inclusions (Eqs. S10-S12). The details of derivations and complete equations are presented in the Supplementary Material.

## Results and Discussion

### Estimation of the radius of induced scission necks by membrane-remodeling proteins from SAXS data

A range of proteins can remodel lipid membranes to form scission necks. The size of scission necks induced by proteins is a key geometric parameter in mediating different processes such as fission, fusion, and membrane trafficking. Here, we use our mechanical framework to estimate the radius of scission necks induced by membrane-remodeling proteins from SAXS measurements and systematically explore the influence of cubic lattice constant, protein area in contact with the membrane, and interfacial tension on the size of the scission necks.

We use the deviatoric curvature (*D*_0_) estimated based on the lattice constants of cubic structures (here we used *Pn3m*) measured in SAXS as input to our model (Fig. 2A). We plotted the change in the system (Eq. 4) from a spherical vesicle to the idealized geometry of the scission neck as a function of the neck radius for four different cubic lattice constants in Fig. 2B. Here, we set the effective surface area of the protein in contact with membrane (*a_p_*) = 4 nm^2^, *S*_0_ = 10 nm and *γ* = 0. Considering an induced *Pn3m* cubic structure, we found that by decreasing the cubic lattice constants, e.g., a = 100 nm to a = 20 nm, the radius of the stable neck at the minimum energy reduces from *r_n_* = 47.52 nm to *r_n_* = 6.75 nm (arrows in Fig. 2B). This implies that proteins that induce cubic phases with smaller lattice constants can strongly bend the membrane and significantly squeeze the scission neck (blue line in Fig. 2B). The size of the scission neck increases almost as a quadratic function of the cubic lattice constant (Fig. S1A). However, for large lattice constants (a > 110 nm), the change in the energy increases monotonically as the neck radius decreases (gray line in Fig. 2B). This suggests that the induced deviatoric curvature by proteins is not strong enough to constrict the membrane and form the scission neck (gray line in Fig. 2B).

The strength of interaction between the embedded proteins and the surrounding membrane depends on the surface area of proteins in contact with the membrane ^50,51^ (*a_p_* in Eq. S16). Proteins with larger contact surface areas bind more strongly to the membrane and can constrict the membrane neck to smaller radii (Fig. 2C, a = 20 nm, *S*_0_ = 10 nm and *γ* = 0). Interestingly, we observed that for proteins with a small contact surface area, an energy barrier is associated with membrane neck constriction (Figs. 2B and 2C). Based on our results, the height of the energy barrier reduces with decreasing cubic lattice constant (Fig. 2B) and vanishes for proteins with large contact surface area (Fig. 2C). These results suggest that changing the effective area of membrane-remodeling proteins in contact with the membrane through oligomerization (such as Drp1 and Dnm1) or the conformational movements can be a driving mechanism that scission machinery exploits to overcome the energy barrier associated with the constriction process of membrane necks ^52^.

In biological contexts, membrane-remodeling proteins have a wide range of sizes and can alter their effective contact area with the membrane. We use our model to construct a contour plot of the induced scission neck for a range of cubic lattice constants and protein surface area in Fig. 2D (*S*_0_ = 10 nm and *γ* = 0). The vesicle remains undeformed and there is no neck in a small domain with a very large cubic lattice constant and a small protein surface area (Fig. 2D). Conversely, narrow necks (*r_n_* < 6 nm) can be formed for small cubic lattice constants (*a* < 16 nm), independent of the protein size (Fig. 2D). We also found that for small cubic lattice constants, the size of the scission neck weakly depends on the interaction between the membrane and the proteins. Thus, to further squeeze the membrane neck and complete the scission phenomena, the induced line tension at the boundaries of protein phase separation can play an important role ^44^ (Fig. 2E, *a_p_* = 4 nm^2^ and *S*_0_ = 10 nm). For example, in the case of *a* = 100 nm, increasing the magnitude of line tension from *γ* = 0 to *γ* = 10 pN ^53,54^ results in an almost 80% decrease (from *r_n_* ~ 47.3 nm to *r_n_* ~ 9.7 nm) in the radius of the scission neck (Fig. 2E). Previous studies have also shown that an increase in the effective line tension, due to the insertion of the M2 transmembrane domain into the membrane, enhances curvature generation and vesicle budding ^55,56^.

Intuitively, one expects that the length of the catenoid (*S*_0_) in the idealized geometry must affect the radius of the scission neck, as it defines the length scale of curvature modulation (Fig. 2F, *a_p_* = 4 nm^2^ and *γ* = 0). In our model, the length of the catenoid has a dual effect on the membrane-inclusion interaction energy (Eq. 1). On the one hand, an increase in the length of the catenoid leads to an increase in the area of the scission neck and the energy cost for membrane constriction. On the other hand, catenoids with larger lengths have smaller deviatoric curvatures 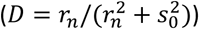, which makes the formation of a narrow neck easier with a lower bending energy cost. The antagonistic effects of *S*_0_ on the elastic mismatch energy cost can be seen in Fig. 2F. At small cubic lattice constants (*a* < 20 nm), the larger surface area of the catenoid becomes dominant in the elastic energy. Therefore, the neck radius increases with increasing the catenoid length (Figs. 2F and S1B). However, at large lattice constants (*a* > 60 nm), the dominant term in the interaction energy is the membrane curvature. Thus, an increase in the catenoid length leads to a smaller neck radius (Figs. 2F and S1C). Interestingly, at intermediate lattice constants (30 nm < *a* < 60 nm), we found that the neck radius is a non-monotonic function of *S*_0_; as *S*_0_ increases, the neck radius decreases and then increases again (Figs. 2F and S1D).

It is possible to construct a dimensionless quantity, the spontaneous scission number (SSN): 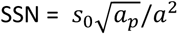 which combines the effects of the cubic lattice constant (*a*), protein surface area (*a_p_*), and catenoid length (*S*_0_). We found that SNN governs the formation of fission necks (Fig. S2). As the SSN increases, the neck becomes smaller, and the radius of the neck is approximately proportional to the power function of the SSN, *r_n_* ~ SSN ^−0.2^ (Fig. S2). Our findings suggested that SSN greater than 1 can give rise to the formation of sufficiently narrow necks (with radii below 5 nm) that allow spontaneous hemi-fission.

### The morphology of membrane caps influences the radius of induced scission necks by membrane-remodeling proteins

Membrane budding is a common phenomenon that often takes place in large vesicles, known as “mother” vesicles, and results in the formation of smaller “daughter” vesicles. This process is observed in various cellular processes, such as clathrin-mediated endocytosis ^57,58^, the formation of synaptic vesicles, and the viral/parasite replication cycle ^59^, where a small coated pit connects to a large mother vesicle via a neck-like structure ^5,60^. To explore the robustness of neck formation by membrane-remodeling proteins in large vesicles, we revised our idealized geometry and assumed that the spherical caps have two different radii, *R* and *R*\* (Fig. 3A). Here, we set *a_p_* = 4 nm^2^, *S*_0_ = 10 nm and *γ* = 0 and plotted the percentage of change in neck radius compared to the case with two identical spherical caps, *R*\*/*R* =1 (Fig. 3A). Based on our results, the scission neck gets wider with increasing ratio of asymmetry between two spherical caps (*R*\*/*R*) (Fig. 3A). The percentage of increase in the neck radius is proportional to the cubic lattice constant (Fig. 3A). For example, for *a* = 20 nm, the neck radius increases less than 2% when *R*\*/*R* =1 increases to *R*\*/*R* = 5. However, our results show that for large lattice constants (*a* = 40 nm), there is a ~10% enlargement in the neck radius as the size of the mother vesicle increase to *R*\*/*R* = 5 (Fig. 3A).

**Figure 3.**
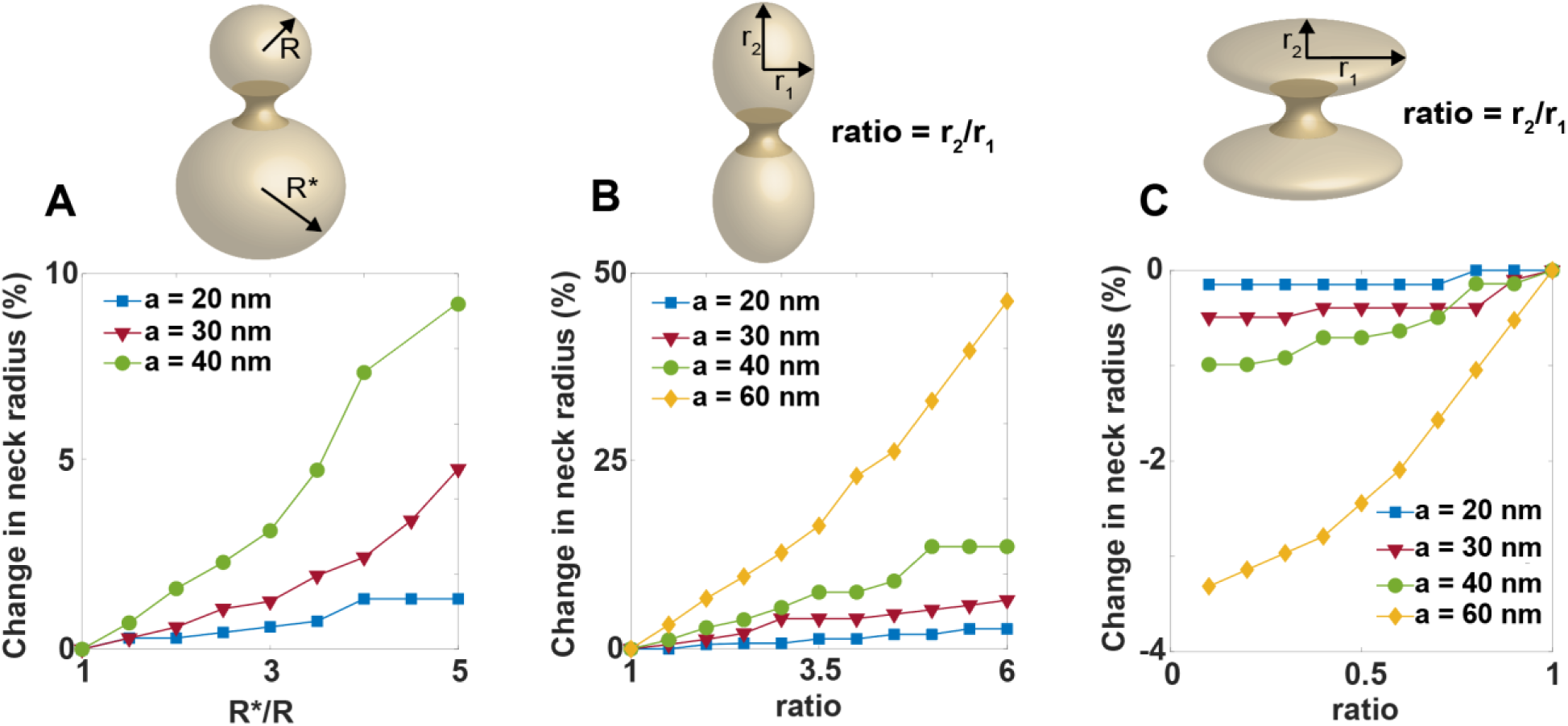
The effects of membrane cap morphology on the radius of induced scission neck by membrane-remodeling proteins. **(A)** Increase in the scission neck radius with increasing size ratio between two spherical caps (R*/R). The scission neck radius exhibits a more pronounced increase as a function of R*/R for larger cubic lattice constants. **(B)** Increase in the scission neck radius with the increasing aspect ratio of the prolate-shaped caps. The size of the neck increases significantly (~50%) when the aspect ratio increases from 1 (spherical caps) to 6 (prolate-shaped caps) at *a* = 60 nm. **(C)** Decrease in the scission neck radius with decreasing aspect ratio of the oblate-shaped caps. For small cubic lattice constants (*a* < 40 nm), the change in the neck radius is negligible (< 1%). In all numerical calculations, we set *a_p_* = 4 nm^2^, *S*_0_ = 10 nm and *γ* = 0.

We also investigated the impact of aspherical membrane cap morphology on the radius of induced scission necks by membrane-remodeling proteins (Figs. 3B and C). This can be a representation of (*I*) the tubular membrane invagination in yeast endocytosis ^61^ and (*II*) the aspherical morphology of the membrane wrapping of elongated viruses such as coronavirus, Ebola, Marburg, and bullet-shaped Rhabdoviruses ^62–66^. Influenza A virus is also known as a pleomorphic virus which displays a range of morphologies from filamentous to spherical ^67^. Here, for simplicity, we only considered prolate-shaped caps with an aspect ratio (*r*_2_/*r*_1_) > 1 (Fig. 3B) and oblate-shaped caps with an aspect ratio < 1 (Fig. 3C).

We found that the size of the scission neck increases with an increase in the aspect ratio of the prolate-shaped cap (Fig. 3B). Particularly, we observed that for large lattice constant (*a* = 60 nm), the radius of the scission neck increases ~50% when increasing the aspect ratio from ratio = 1 to ratio = 6 (Fig. 3B). For oblate-shaped caps, our results show that the neck radius slightly decreases (< 4%) when decreasing the aspect ratio from ratio = 1 to ratio = 0.1 (Fig. 3C). However, both prolate and oblate-shaped caps create a large energy barrier in neck constriction (Figs. S3 and S4). This could be related to the high membrane bending energy in ellipsoidal caps compared to the spherical caps ^68^. We observed that the height of the energy barrier increases for larger lattice constants and is more dramatic in oblate-shaped caps (Figs. S3 and S4). Overall, these results suggest that the process of membrane budding from larger vesicles and with an aspherical cap morphology requires a higher degree of curvature induction by membrane-remodeling proteins.

### Scission neck from viral budding protein is smaller than that from mitochondrial fission proteins, and exhibits different lipid requirements

It is interesting to test the model by comparing the behavior of induced scission necks by viral budding M2 protein from influenza A virus versus dynamin-mediated mitochondrial fission in eukaryotic cells. We have previously shown that Dnm1 and the M2 protein from influenza A virus can remodel lipid membranes to cubic structures with NGC ^22,23^. Using these SAXS measurements as inputs for our model and estimating the effective surface area of proteins based on their X-ray crystallography, we calculated the radius of constricted necks induced by M2 protein and Dnm1 in different lipid compositions (Fig. 4A). In these calculations, we considered 20% variations in the protein surface area (*a_p_* ± 20%) and the length of neck (*S*_0_ ± 20%), assuming no line tension, *γ* = 0 (Fig. 4).

**Figure 4.**
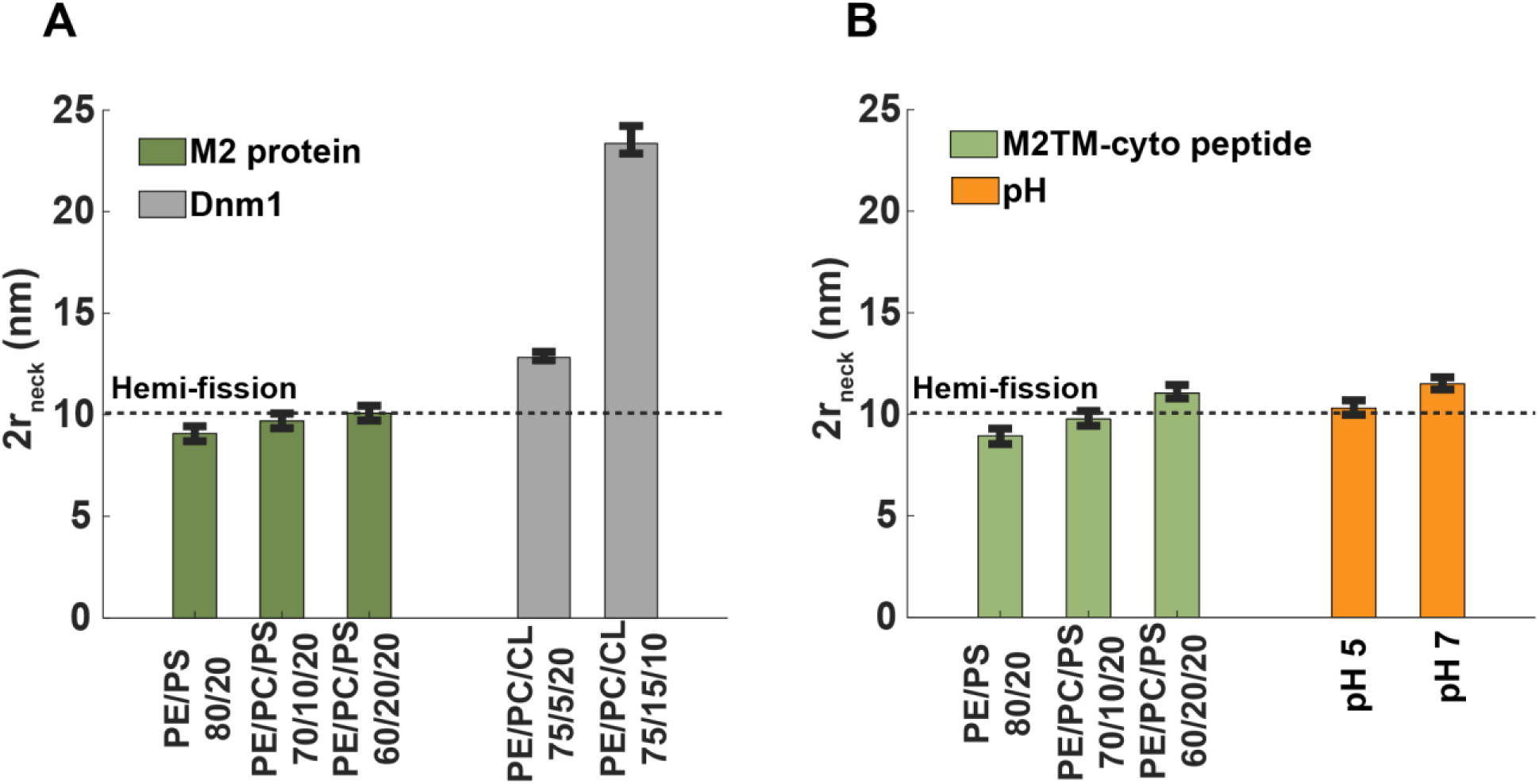
Comparison of membrane remodeling activity and induced scission necks by viral protein M2 from influenzas A and eukaryotic fission protein Dnm1. **(A)** Estimation of induced scission necks by M2 protein of influenza A virus and Dnm1 in different lipid compositions, using our mechanical framework and the cubic lattice constants measured in our previous SAXS experiments ^22,23^. The M2 protein of influenza A virus can form narrow constricted necks < 10 nm within the range of possible hemi-fission regime in different lipid compositions. **(B)** Estimation of induced scission necks by M2TM-cyto peptide in different lipid compositions, and the full length M2 protein under neutral (pH 7) and acidic (pH 5) conditions. The M2 induces smaller scission neck sizes in acidic conditions. In panels (C) and (D), we considered 20% variations in neck length (*S*_0_ = 10 nm) and effective contact surface area *a_p_* (*γ* = 0).

Based on our calculations, the size of induced fission necks by Dnm1 highly depends on the percentage of conical cardiolipin lipids present in the membrane (Fig. 4A). For instance, in model membranes with ternary phospholipid mixtures of phosphatidylethanolamine (PE), phosphatidylcholine (PC), and cardiolipin (CL) at molar ratios of 75/5/20 and 75/15/10 PE/PC/CL, Dnm1 can constrict the membrane into catenoid-shaped necks with diameters ranging from 10.78-11.44 nm and 22.92-23.4 nm, respectively (Fig. 4A). Previous studies have also shown that membrane lipid composition plays an important role in mitochondrial fission ^69–71^. In particular, the presence of conical lipids in the mitochondrial membrane facilitates the interaction and activation of the GTPase domain of fission proteins ^69–71^. Using a previously proposed catenoid-shaped neck model ^30^, the average amount of Gaussian curvature in a cubic phase induced by Dnm1 in 75/5/20 PE/PC/CL model membrane maps to a fission neck of approximately 12.6 nm in diameter ^22^. This is in agreement with our estimated neck size of 10.8 nm < 2*r_n_* < 11.5 nm using the dumbbell shaped constricted neck in our model (Fig. 4A). In this context, it is interesting to note that connecting the catenoid-shaped neck to spherical caps with positive Gaussian curvature can decrease the size of the neck.

We employed our mechanical framework to estimate the size of the pinching neck generated by M2 proteins under different conditions; *(I)* the full length of M2 protein of the influenza A virus in different lipid membrane compositions ^23^ (Fig. 4A), *(II)* a synthesized M2TM-cyto peptide in various lipid membrane compositions ^23^ (Fig. 4B), *(III)* the full length of M2 protein in both neutral (pH 7) and acidic (pH 5) buffer conditions (Fig. 4B). Here, we assume that the viral budding occurs from a large vesicle, with a relative size ratio of *R*\*/*R* = 5. We found that both full length M2 protein and the synthesized M2TM-cyto peptide can form narrow necks (neck diameter < 10 nm) which can trigger the spontaneous membrane hemi-fission reaction ^72^ (Figs. 4A and 4B). Interestingly, we found that, unlike fission proteins, the diameter of the endocytic neck generated by the M2 protein and the synthesized M2TM-cyto peptide does not strongly depend on the lipid composition (Figs. 4A and 4B). This is interesting for several reasons. That viruses are much smaller than mitochondria places more stringent requirements for scission. Moreover, viruses, in principle, need to remodel host membranes with a wide range of lipid compositions.

The M2 viroporin is a proton pump as well as a membrane remodeling protein. We observed that under acidic conditions, the M2 protein from influenzas A induces tighter NGC and can constrict endocytic necks to narrow sizes falls within the range of the spontaneous hemi-fission regime (Fig. 4B). This indicates that the two functions of M2 both promote budding and do not act against one another. This could be attributed to a change in the backbone conformation of the C-terminal helices, which leads to altered lipid-protein interactions and induces anisotropic curvature, similar to the opening and activation of the influenza A M2 proton channel in response to acidification ^35^.

### M2 protein from the influenza A virus can robustly induce anisotropic curvature in a range of lipid compositions in coarse-grained molecular dynamics simulations

To test the predictions from the theoretical framework, we performed coarse-grained molecular dynamics simulation to compare the capability of the M2 protein of the influenza A virus versus Dnm1 to induce anisotropic curvature on flat membranes with two different lipid compositions (Fig. 5). Previous studies have shown that the M2 proton channel and the pleckstrin homology (PH) domain of Dnm1 contains amphiphile loops that can insert into the lipid membrane and constrict the neck ^23,73–76^. Using a previously trained support vector machine (SVM) classifier to assess the membrane-restructuring activity of a given protein sequence ^77^, we showed that both the M2 proton channel and the PH domain of Dnm1 contain amino acid sequences capable of remodeling membranes through anisotropic membrane curvature induction (Fig. S5).

**Figure 5.**
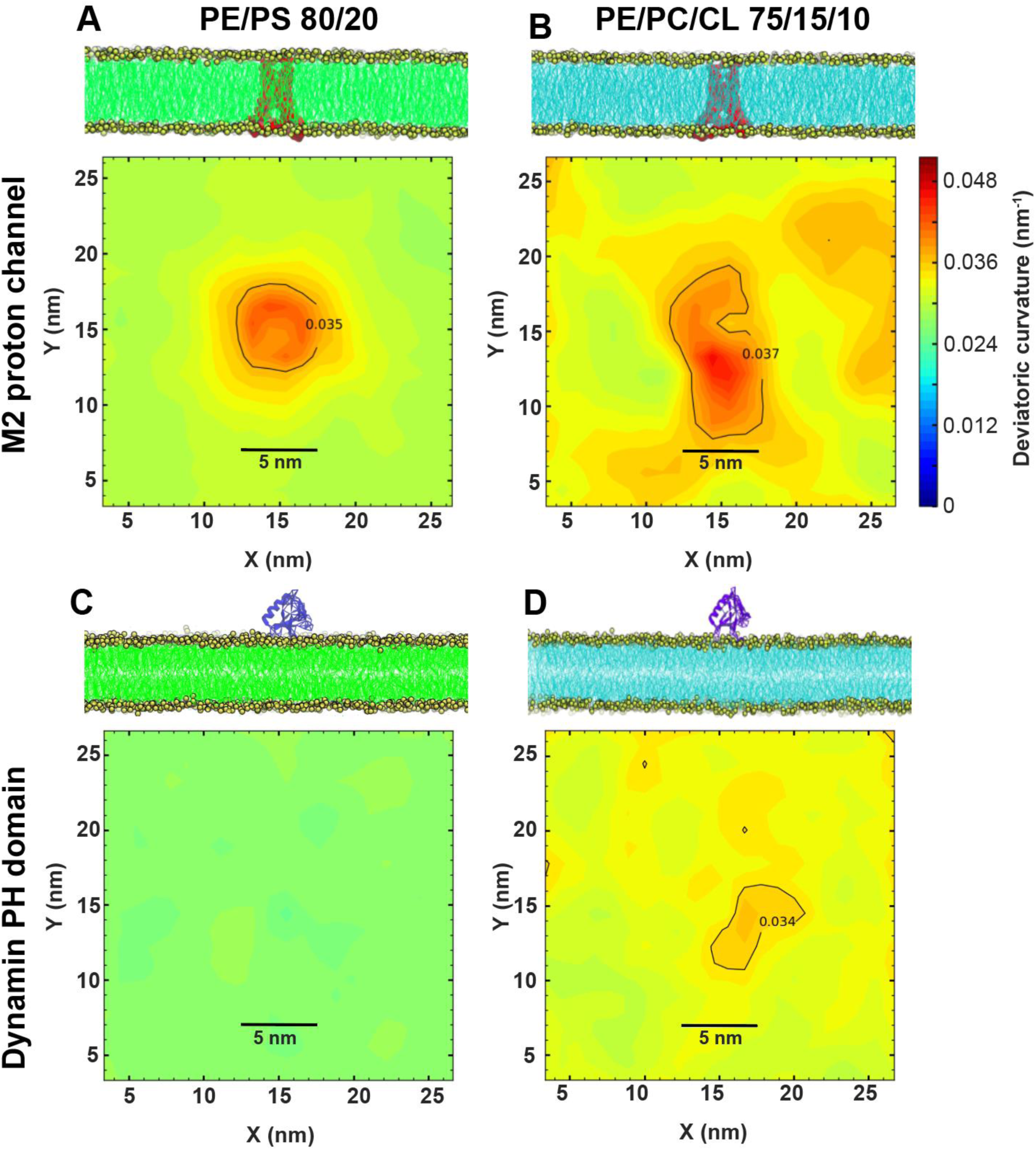
M2 protein from the influenza A virus induces robust anisotropic curvature independent of the membrane lipid composition in molecular dynamics simulations. Structure of the protein-membrane complex (upper panel) and contour plot of the deviatoric curvature generated by the protein on the membrane surface (lower panel) for **(A)** M2 proton channel from the influenza A virus on PE/PS 80/20 lipid bilayer, **(B)** M2 proton channel from the influenza A virus on PE/PC/CL 75/15/10 lipid bilayer, **(C)** Dynamin PH domain on 80/20 PE/PS lipid bilayer, and **(D)** Dynamin PH domain on PE/PC/CL 75/15/10 lipid bilayer. M2 protein and dynamin PH domain are shown by the red and violet colors, respectively.

We placed a single transmembrane M2 proton channel at the center of a flat lipid bilayer with two different lipid compositions: (I) 80/20 PE/PS and (II) 75/15/10 PE/PC/CL (Figs. 5A and 5B). The system equilibrated for 2μs and we used the equilibrated trajectory to compute the deviatoric curvature generated on the membrane by the protein. The deviatoric curvature was calculated using the MDAnalysis tool and the VTK library of Python ^78^. Details of the simulation protocol are described in the Supplementary Material. We found that M2 protein can generate a similar degree of anisotropic curvature on both lipid compositions (Figs. 5A and 5B). However, our results show that the induced anisotropic curvature in 75/15/10 PE/PC/CL lipid composition extends to a relatively larger distance compared to the 80/20 PE/PS flat lipid bilayer, which is likely due to the presence of flexible cardiolipins (Figs. 5A and 5B). These simulation results are in agreement with predictions of our theoretical model and SAXS measurements that M2 protein can robustly induce deviatoric curvature on membranes with different lipid compositions.

As expected, deviatoric curvature generated by the PH domain of Dnm1 strongly depends on the lipid composition of the membrane. While it induces a decent amount of deviatoric curvature on the 75/15/10 PE/PC/CL membrane, it does not generate any deviatoric curvature near the location of the protein on the 80/20 PE/PS membrane (Figs. 5C and 5D). We also observed that the magnitude of the induced deviatoric curvature by the M2 protein is higher than the PH domain on both lipid compositions (Fig. 5). The curvature generated by the M2 protein also extends to a relatively larger distance compared to the induced curvature by the PH domain. These results suggest that neck formation by the dynamin-related family of proteins strongly depends on the lipid composition, while the M2 protein from the influenza A virus can form narrow scission necks over a range of cell membranes with different lipid compositions.

In summary, we present a minimal mechanical model to directly use SAXS measurements to estimate the radius of scission necks induced by membrane-remodeling proteins. Using this method, it is possible for the first time to systematically explore the interplay between the protein surface area, the magnitude of induced curvatures by proteins, interfacial tension at the phase separated protein boundary, and the morphology of the caps in regulating the size of scission necks (Figs. 2 and 3). In particular, we introduce a dimensionless quantity, the 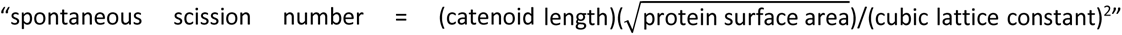 which characterizes the extent of scission neck constriction (Fig. 2). Our results also show that the neck radius depends on the morphology and symmetry of caps (Fig. 3). For instance, for large lattice constants, the neck radius increases noticeably for membrane budding from a large vesicle (~10%) and with prolate-shaped caps (~50%) (Fig. 3). Additionally, we find that for ellipsoidal-shaped caps, neck constriction is associated with a large energy barrier (~1500*kT*) (Figs. S3 and S4). This substantial energy barrier implies that a secondary mechanism such as actin-mediated pinching forces or helical twist is required to complete fission ^61,79,80^.

In the general context of required NGC for membrane remodeling in scission necks versus transmembrane pore formation, we observe that the energy barrier associated with the formation of transmembrane pores ^47,81^ is almost 6x larger compared to the energy barrier involved in the neck constriction process (identical spherical caps in Fig. 2). Additionally, we identify that scission necks can be formed across a wide range of cubic lattice constants, 0 < *a* < 100 nm (Fig. 2), whereas pore formation is confined to a relatively narrow range of lattice constants (0 < *a* < 30 nm) ^47^. Thus, we predict that the membrane remodeling required for scission neck formation is simpler than the initiation of transmembrane pore opening. We find that the proton pumping activity of M2 viroporin does not interfere with its membrane remodeling activity. In fact, the two activities appear to be synergistic. It would be interesting to investigate other pH-dependent microbial pore-formers, such as Diphtheria toxin ^82^. The method presented here is potentially applicable to other protein-induced membrane-remodeling processes such as budding, blebbing, and tubulation, and can provide insight into the mechanical design principles for robust vesiculation, e.g. in drug or gene delivery systems.

## Methods

The complete derivations with details are given in the Supporting Material.

## Supporting information

Complete derivations with details; Supplemental Figure S1 including the scission neck size as a function of cubic lattice constant and catenoid length; Figure S2 shows the variation of a neck radius as a function of the spontaneous scission number; Supplemental Figures S3 and S4 including the change in the energy of the system from a spherical vesicle to the idealized scission neck geometry with prolate and oblate-shaped caps. Supplemental Figure S5 includes the heatmaps of σ scores from an SVM classifier for the M2 proton channel and Dnm1 PH domain.

## Acknowledgments

This work was supported by the American Heart Association (AHA 966662) grant to G.C.L.W. H.A was supported by the Vascular Biology Training Grant (T32 HL069766-21). J.D.A. is supported by the NSF Graduate Research Fellowship Program (DGE-1650604). T.M gratefully acknowledges the support from the Government of India: Science and Engineering Research Board via Sanction No. SRG/2022/000548. We thank the Stanford Synchrotron Radiation Lightsource (SSRL) (Menlo Park, CA, USA) for access to beamline 4-2. Use of the SSRL, SLAC National Accelerator Laboratory, is supported by the U.S. Department of Energy, Office of Science, Office of Basic Energy Sciences under contract no. DE-AC02-76SF00515. The SSRL Structural Molecular Biology Program is supported by the U.S. Department of Energy, Office of Biological and Environmental Research, and by the National Institutes of Health, National Institute of General Medical Sciences (including P30GM133894). T.M. is grateful for the computational resources provided by PARAM Sanganak under the National Supercomputing Mission, Government of India, at the Indian Institute of Technology, Kanpur.

## Author contributions

H.A designed the research and G.C.L.W supervised the research. T.M and S.G conducted the molecular dynamics simulations. H.A, E.W.C.L, J.D.A, R. Y, and G.C.L.W wrote the manuscript. All authors reviewed and approved the final version of the manuscript.

